# Chromosome-level genome assembly for Sichuan taimen (*hucho bleekeri*) reveals the extraordinary tandem repeat proportions and its persistent population shrinkage

**DOI:** 10.1101/2023.12.15.570915

**Authors:** Xinmiao Zhang, Dongmei Xiong, Shenglong Jian, Yu Jiang, Lixin Wang

**Author notes:** Correspondence authors. (Lixin Wnag) and (Yu Jiang).

## Abstract

Salmonid fishes are globally renowned and valuable, with most members of the Salmonidae family living in seawater and exhibiting migratory behavior. In contrast, huchonid fishes (*Hucho* spp.and *Brachymystax* spp.), an ancient evolutionary branch within Salmoninae, are entirely landlocked. The Sichuan taimen (*Hucho bleekeri* Kimura) is a critically endangered fish that has attracted widespread concern and is one of eleven national first-level protected fishes in China. However, genome resources for all *Hucho* spp., including *H. bleekeri*, are scarce, and the genomic characteristics of this ancient evolutionary lineage remain unclear, hindering conservation biology efforts. Here, we present the first chromosome-level genome for the Sichuan taimen, with a final genome size of approximately 3.45 Gb across 44 chromosomes. The Sichuan taimen genome contains 44.15% tandem repetitive sequences, exceeding those of all sequenced salmonid fishes. We also identified 44 Ss4R homeologous block pairs in the assembled genome. Genome synteny analysis suggested a ∼5 fold tandem repeat expansion in the Sichuan taimen compared to the Salmonidae ancestor Northern pike. Phylogenetic analysis estimated the divergence time between huchonid and other Salmoninae fishes at approximately 33.29 million years ago (Mya). The divergence time between Siberia taimen and Sichuan taimen was estimated at around 2.29 Mya, with their effective population size declining from around 1 Mya. The genomic resource provided in this article will promote the protection of the Sichuan taimen and evolutionary genetics studies of salmonids.

## INTRODUCTION

The Sichuan taimen (*Hucho bleekeri* Kimura, 1934) belonging to the subfamily Salmoninae, is a glacial remnant species and endemic to China (Wang et al., 2016). It has the lowest distribution latitude among all taimen species, and is primarily found in the upper tributaries of the Yangtze River and Wei River (Hu et al., 2008). Sichuan taimen, characterized by their large body size and predatory behavior, occupy the top of the aquatic food chain (Wang et al., 2016). However, their late sexual maturation makes them particularly vulnerable to human activities and ecological changes (Allendorf & Hard, 2009; Sadovy et al., 2013). Individuals of Sichuan taimen are found sporadically and scattered in their distribution, and have been listed in China red data book of endangered animals since 1998. Despite this, they continues to be illegally captured by poachers for their high nutritional value and taste (Sung et al., 1998). Due to the continuous decline of the population size, the Sichuan taimen was listed as critically endangered (CR) level in the International Union for Conservation of Nature (IUCN) Red List of Threatened Species in 2012 (Song, 2012). As of 2021, the Sichuan taimen is under the highest level of protection from the Chinese government, and is the only salmonid fish with national protected animal in “class I”. Genomic data provide significant opportunities for advancements in the conservation and evolutionary research of threatened animals (Allendorf et al., 2010; Kohn et al., 2006). As sequencing costs have decreased, the genomes of many endangered fish species have been published, including *Trachidermus fasciatus* Heckel (Xue et al., 2022), *Myxocyprinus asiaticus* (Liu et al., 2022), and *Brachymystax tsinlingensis* (Zhu et al., 2022), etc. Based on genome data, scientific and rational conservation policies have been formulated, enhancing the protection benefits of these endangered animals (Brandies et al., 2019; Waples et al., 2020). However, the lack of genome data for *Hucho bleekeri* creates a gap in nuclear-based phylogenetic research of Salmonidae and affects genomic studies, such as taxonomic studies based on SNPs (Chen et al., 2020; Zhang et al., 2020) and researches on transcriptome (Chen et al., 2021).

Due to the whole genome duplication and lineage-specific rediploidization, the genome of salmonids have became complex and variable, increasing the difficulty of assembly (Dysin et al., 2022; Robertson et al., 2017). The circular consensus sequencing (CCS) mode in single-molecule real-time (SMRT) sequencing by Pacific Biosciences can produce long and high fidelity (HiFi) reads, which facilitates the genome assembly process and improves quality (Wenger et al., 2019). In this research, we utilized HiFi and Hi-C sequencing technology to construct a high-quality genome for Sichuan taimen. From the genome we identified a high proportion of tandem repetitive sequences. Additionally, we analyzed the phylogenetic relationships of salmonid fishes and the divergence history of *H. bleekeri* and *H. taimen*, another species found in Heilongjiang. Our work provides insights into the genome evolution of salmonid fishes and the conservation of the endangered Sichuan taimen.

## MATERIALS AND METHODS

### Sample collection and sequencing

Two wild individuals of *H. bleekeri* were collected from Taibai river (Baoji, Shaanxi province), one for genome assembly the other for resequencing. Additionally, a cultured individual of *H. taimen* was collected from Heilongjiang. All the experiments in this study were performed in accordance with the guidelines of China Council on Animal Care, and approved by the Experimental Animal Ethics Committee of Northwest A&F University (NWAFU-314020038).

Genomic DNA was extracted from kidney tissue with cetyltrimethylammonium bromide (CATB) method (Stefanova et al., 2013). The reads for genome size estimation were generated on Illumina NovaSeq-6000 platform after the constructed of 150 bp library. For genome assembly, a 15 kb library preparation was performed by SMRTbell Express Template Prep Kit 2.0 (Pacific Biosciences, CA, USA), and sequenced on the PacBio Sequel II platform (Pacific Biosciences, CA, USA) with two cells of single molecule real time sequencing (SMRT). The CCS software v.6.0.0 (https://github.com/PacificBiosciences/ccs) was performed to integrate HiFi reads with the parameters “--min-passes 3 --min-length 10 --min-rq 0.99”. The Hi-C library was constructed and sequenced on Illumina NovaSeq-6000 platform. The RNA samples were extracted from fin, brain, gill, gonad, and eye, then sequenced on Illumina NovaSeq-6000 platform. The two wild samples for demographic history analysis were sequenced on DNB-seq platform (BGI).

### Genome size estimation

The raw Illumina paired-end reads of Sichuan taimen were filtered using Fastp version 0.20.0 (Chen et al., 2018) with default parameters. The genome size estimation of Sichuan taimen was performed by GCE software version 1.0.2 (Liu et al., 2013). A total of 58.89 Gb clean reads were used to calculate the distribution of k-mer frequency using kmerfreq with k=17. Then the gce software was ran in both homozygous and heterzygous mode to evaluate the genome size and heterozygosity. The genome heterozygous was calculated using the formula “a[1/2]/k”.

### Chromosome level genome assembly

The PacBio HiFi reads were assembled using Hifiasm version 0.13-r308 (Cheng et al., 2021) to obtain *de novo* contigs. After quality control by Fastp (Chen et al., 2018), the clean Hi-C reads were used to integrate the contigs. The Hi-C interaction matrix was generated using Juicer (version 1.6)(Durand et al., 2016b)scripts with -s MboI parameter, then, the 3D-DNA (version 201008)(Dudchenko et al., 2017)with 0 rounds was used to link the contigs into chromosome level based on the Hi-C interaction matrix. The initial assembly was manually corrected according to the Hi-C heatmap using Juicebox version 1.11.08 (Durand et al., 2016a) and run the 3D-DNA again to generate the final assembly. The BUSCO (Benchmarking universal single-copy orthologues) version 3.0.2 (Simão et al., 2015) was used to assess the accuracy and integrity of the genome with the gene database of ray-finned fishes (actinopterygii_odb9).

### Repeat sequence detection and gene annotation

To recognize repeat sequences of the Sichuan taimen genome, we build a *de novo* library of transposable elements (TEs). First, the RepeatModeler version 2.1 (Flynn et al., 2020) with default parameters was ran to constructed a repeat library; then an additional long terminal repeat (LTR) library of the Sichuan taimen was built by LTR_FINDER version 1.07 (Xu & Wang, 2007) and LTR_RETRIEVER version 2.9.6 (Ou & Jiang, 2018); third the final repeat library was integrated by replacing the LTR elements of the first repeat library with the LTR library. To further classify the “Unknown” sequence, the BLAST version 2.14.0 (Ye et al., 2006) was performed between the final repeat library and public repeat library (http://www.hrt.msu.edu/uploads/535/78637/Tpases020812.gz).

The TEs of the genome were described and masked by RepeatMasker version 4.1.2 (Tarailo-Graovac & Chen, 2009) with parameters “-xsmall -s -cutoff 255 -frag 20000 –gff -engine ncbi”. The tandem repeats (TRs) were detected through Tandem Repeats Finder (TRF) version 4.09 (Benson, 1999) with parameters “2 7 7 80 10 50 500 -f -d -m”. The TRs were further divided into short tandem repeats (STRs) and variable number tandem repeats (VNTRs) according to the length of repeat unit (STR: unit number at 1 to 6; VNTR: unit number more than 6).

The gene annotation of the Sichuan taimen genome was carried out using BRAKER2 pipeline version 2.1.5 (Hoff et al., 2016; Hoff et al., 2019). For transcriptome based prediction, the RNAs were mapping to the assembled genome using STAR version 2.7.1a (Dobin et al., 2013) with default parameters, then the RNA sequence data were used to predict genes through auto BRAKER2 pipeline. For homology based prediction, the protein sequence of six species including *Lepisosteus oculatus* (GCF_000242695.1)(Braasch et al., 2016), *Esox Lucius* (GCF_011004845.1)(Ishiguro et al., 2003), *Oncorhynchus mykiss* (GCF_013265735.2)(Gao et al., 2021), *Oncorhynchus nerka* (GCF_006149115.2), *Salmo salar* (GCF_000233375.1)(Lien et al., 2016), and *Brachymystax tsinlingensis* (GCA_031305455.1) which represent the sister clade or the ancestor species of taimen were download, and predicted through BRAKER2 pipeline. The results from RNA and protein-based annotation were combined using TSEBRA version 1.0.3 (Gabriel et al., 2021) with default parameters to generate the final gene prediction file. The gene expression level was measured by transcript per million (TPM) which were calculated using Kallisto (Bray et al., 2016) with default parameters. To remove the pseudogenes caused by tandem repetitive sequence, the gene set was filtered based on the following criteria: 1) contain only one exon; 2) TPM<1; 3) no matched BLAST results with *B. tsinlingensis*.

### Chromosome synteny identification

Homology blocks were reconstructed using whole-genome duplicated genes. Diamond (v0.8.24.86) with default parameter was used to align against the predicted genes. Then selected paired genes with highest identity and identity ≥ 75% from the Diamond results. Finally, one-to-one paralogs were extracted and conducted manual linkage. The *Esox lucius* genome was download from NCBI (GCF_011004845.1).

### Phylogenetic analysis

Protein sequences of 10 species *Esox lucius*, *Coregonus* sp. *’balchen’* (GCA_902810595.1)(Christensen et al., 2020), *Thymallus thymallus* (GCA_023634145.1), *Oncorhynchus mykiss*, *Oncorhynchus nerka*, *Salvelinus namaycush* (GCF_016432855.1)(Smith et al., 2022), *Salmo salar*, *Salmo trutta* (GCF_901001165.1)(Hansen et al., 2021), *Hucho bleekeri*, and *Brachymystax tsinlingensis* were used to construct phylogenetic tree. The single copy orthologous proteins among the 10 species were recognized by Orthfinder version 2.3.12 (Emms & Kelly, 2019) with “msa” method. After alignment using MUSCLE version 3.8.31 (Edgar, 2004), the species tree was calculated through RAxML version 8.2.12 (Stamatakis, 2014) with 1000 bootstraps. The expansions and contractions of gene family was estimated using CAFE5 (Mendes et al., 2020).

The divergence time among all these species were calculated using MCMCtree program of PAML software version 4.9d (Yang, 2007). Four fossil records were downloaded from TimeTree (Kumar et al., 2022) and calibrated the results, the Markov chain Monte Carlo (MCMC) was sampled with every 1,000 step and conduct 10,000 samples, the pre-burnin number was set at 20,000.

### Divergence time estimation of two taimen species

The raw resequencing data of *H. taimen* was mapping to the assembled genome using BWA-MEM version 0.7.17 after quality control. The whole genome sequence for *H. taimen* was generated using angsd version 0.918. To obtain the gene annotation information of *H. taimen*, the liftoff (v1.6.3) was operated with default parameters, and further removed genes with a proportion of unknown nucleotides above 1%. Then, four species were selected to generated 1:1 single copy genes using blastn version 2.2.31+, including *H. taimen*, *H. bleekeri*, *B. tsinlingensis*, and the outgroup *Salmo salar*. After the construction of phylogenetic tree using the methods described above, we estimated the divergence time using BEAST version 2.6.60, the nucleotide substitution model F81 was predicted by jmodeltest version 2.1.10, the chain length was set to 50,000,000 and pre-burnin 10%.

### Demographic history estimation

The pairwise sequentially Markovian coalescent (PSMC) method version 0.6.5-r67 (Li & Durbin, 2011) was utilized to predict the effective population size. Besides the two wild taimen sample, we used two lenok samples with highest sequencing depths from Heilongjiang River (*B.tumernsis*) and Qinling Mountains (*B. tsinlingensis*). Due to the different sequencing depth, we used different strategies to generate the diploid consensus sequences for taimen and lenok, -D 60 for taimen and -D 20 for lenok at vcfutils.pl packages of bcftools version 1.12 (Danecek et al., 2021). The psmc was run with parameters “-N25 -t30 -r4 -p 8+4+25*2+4+6” to control the divergence at left of the results.

## RESULTS

### Genome size estimation and assembly

A total of 426,685,475 paired-end reads were obtained from the Illumina NovaSeq-6000 platform, yielding a total length of 64.00 Gb. The genome size estimated by GCE (k=17) was ∼3.21 Gb with a low heterozygous ratio of 0.44% (Supplementary Figure S1; Supplementary Table S1). Based on the estimated genome size, the PacBio HiFi and Illumina Hi-C sequencing produced 76.48 Gb (23.8×) and 350.63 Gb (109.23×) of raw reads for genome assembly, respectively. Besides, 155.88 Gb of resequencing data from two additional wild samples (∼22 ×) were generated. After *de novo* genome assembly using the HiFi reads, a 3.48 Gb primary genome was generated, which included 4,373 high quality contigs (>2 Kb) with a contig N50 of 2.4 Mb (Table 1). Following Hi-C assisted genome assembly and scaffold deduplication, the contigs were anchored into 44 chromosome-level scaffolds, covering 95.26% of the total scaffolds length, with a scaffold N50 of 73.71 Mb (Figure 1A; Supplementary Figure S2; Table 1). The final Sichuan taimen genome, closely matching the estimated size, had a total size of 3.45 Gb with a GC content of 41.99% (Figure 1A, B; Table 1). The BUSCO results showed that the assembled genome included 95.3% complete genes from the actinopterygii_odb9 database, demonstrating high completeness (Supplementary Table S2).

**Figure 1.**
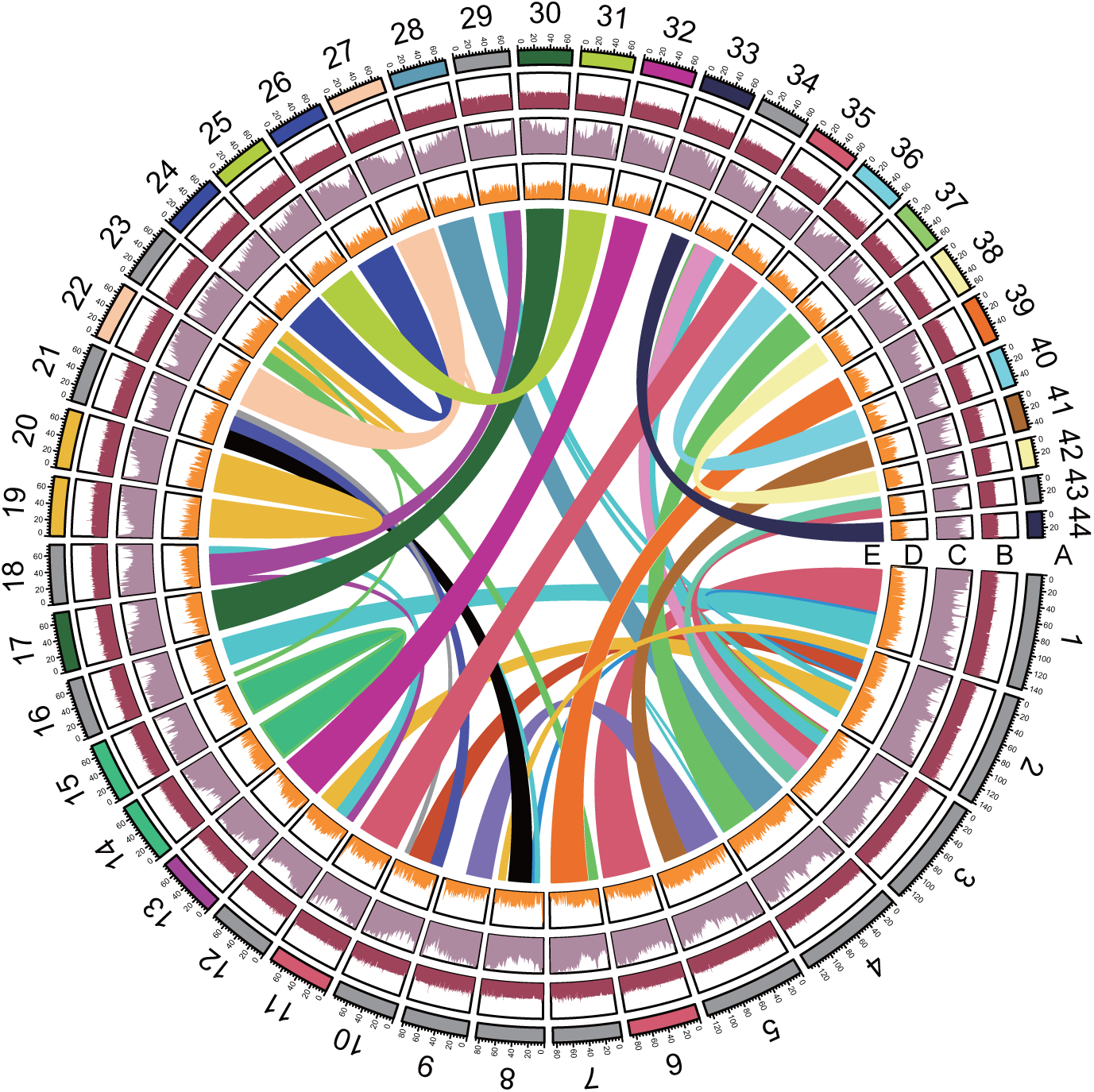
Circos plot of the genome assembly statistics of Sichuan taimen. A: Chromosome names and length scales. Chromosomes consisting of multiple homologous regions were shown in gray (The same below). B: GC ratio (scale: 0-100%). C: Proportion of total repeats (TR and TE)(scale: 0-100%). D: Gene density after log transformation (scale: 0-7). E: Homeologous regions in the Sichuan taimen genome recognized by 1:1 duplicated proteins.

**Table 1.** Summary statistics of the Sichuan taimen genome assembly.

| Category | Length (bp) |  | Number |  |
| --- | --- | --- | --- | --- |
|  | Contig | Scaffold | Contig | Scaffold |
| Max | 25,857,850 | 143,121,144 | - | - |
| Min | 3,656 | 18,648 | - | - |
| N50 | 2,400,826 | 73,705,547 | 354 | 19 |
| N90 | 418,871 | 59,067,955 | 1680 | 39 |
| Total | 3,479,724,502 | 3,451,243,059 | 4373 | 505 |

### Landscape of repeat sequences

The non-redundancy length of the total repeat sequences recognized by three annotation methods was 2.60 Gb, covering 74.21% of the assembled genome, which represents the highest proportion of repetitive sequences among salmonids (Figure 1C; Figure 2; Supplementary Table S3, S4 and S5). The interspersed repeats reported by the Repeatmasker and LTR_FINDER composed 36.18% of the genome, the lowest proportion among salmonids; however, the total length of interspersed repeats in the Sichuan taimen genome was similar to that of other salmonids (Supplementary Table S3 and S4). The proportion of retroelements in the Sichuan taimen genome was 13.11%, with 7.03% LINEs and 5.98% LTR elements (Supplementary Table S3). DNA transposons composed 9.93% of the genome, with the Tc1-IS630-Pogo family being the most abundant TE, similar to other salmonids, covering 9.38% of the Sichuan taimen genome (Supplementary Table S3).

**Figure 2.**
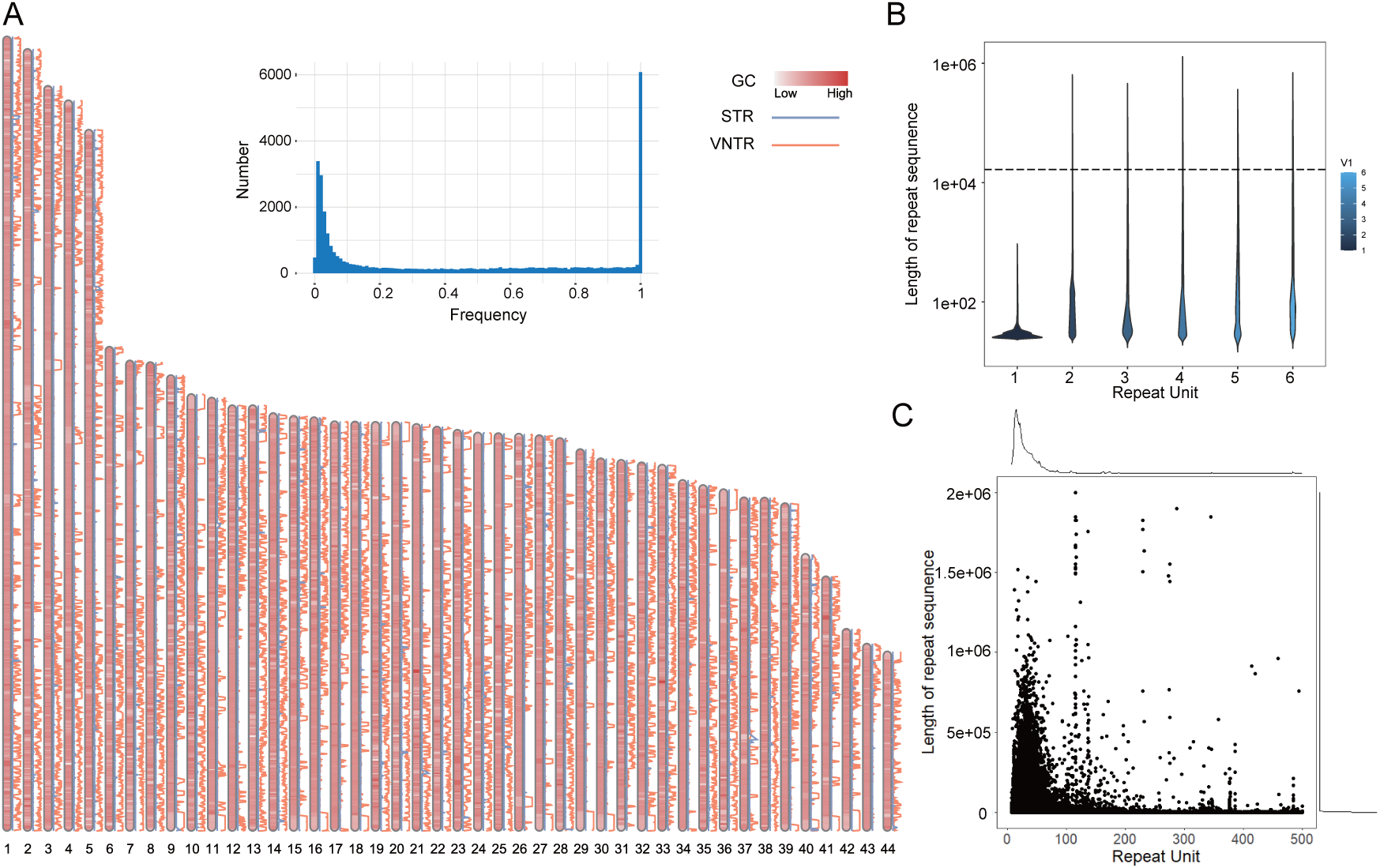
Distribution and statistics of repetitive sequence in the Sichuan taimen genome. A: Visualization of genomic GC content, STR, and VNTR distribution based on 100 Kb windows; the bar graph reflects the TR frequency distribution in genome-wide 100 Kb windows. B and C were statistics of tandem repeat unit length and total tandem repeat sequence length of STRs and VNTRs respectively; The average length of the HiFi reads (16620 bp) was labeled as dashed line.

Tandem repetitive sequences covered 44.15% of the Sichuan taimen genome, representing highest proportion among salmonids (Figure 2A; Supplementary Table S5). The lengths of STRs and VNTRs were 45,561,499 bp and 1,553,872,291, accounting for 1.30% and 44.40% of the genome, respectively. Additionally, 59.12% of STRs were less than 1 Kb away from the surrounding VNTRs (Figure 2A). The distribution of TR was not uniform. We found that 29.44% of the regions in the Sichuan taimen genome had low TR ratios (One-tenth of the genome-wide scale; Figure 2A), and some chromosomes had an enrichment of VNTR at their ends (Figure 2A). The distribution of STRs in the Sichuan taimen genome was relatively scattered, with only the top 200 STRs by length exceeding the average length (16,620 bp) of the HiFi reads (Figure 2B). Dinucleotide repeats accounted for 81.53% of STRs (Supplementary Table S6), which is higher than the average level for vertebrates (Adams et al., 2016). We identified 3,337 VNTRs with lengths over 100 Kb, totaling 0.97 Gb and covering 28.20% of the genome (Figure 2C). In addition, the presence of TRs affected GC content in local regions of the genome; however, the GC content of the TR regions was 41.16%, similar to the genome-wide range (Figure 2A).

### Gene annotation and genomic synteny

A total of 40.00 Gb of RNA reads were generated from five tissues for RNA-based gene annotation. After filtering pseudogenes, the BRAKER2 annotation pipeline predicted 49,052 genes, with an average length of 14,577 bp, containing 338,448 exons in total. The average number of exons per gene was ∼7 (Figure 1D). After mapping to the NR databases, only 79.02% of the predicted genes were functionally annotated. The TR ratio in intron regions was 22.74%, significantly lower than the genome-wide level (Z-test; P<0.01). Based on kernel density estimation, 161 gene hotspots were identified (Supplementary Figure S3). NR database annotation results showed that the top three regions with the highest gene density were located in histone gene family loci on chromosome 8 and 22. The BUSCO analysis for the predicted genes indicated that 88.1% were complete BUSCOs (C: 88.1%[S: 43.3%, D: 44.8%], F: 4.3%, M: 7.6%, n: 4584).

After the protein mapping process, we identified 15,787 paired homeologous genes, accounting for 64.37% of the total predicted genes. Eventually, 44 homeologous blocks were located based on the genomic synteny of the genes (Figure 1E; Supplementary Supplementary Table S7). The circos plot of homeologous blocks showed that the longer chromosomes were more likely to be composed of multiple homologous blocks, especially for the first five chromosomes larger than 120 Mb (Figure 1E). Notably, more than half of the chromosomes (27/44) were composed of a single homologous block, and the repeat distribution pattern was similar between blocks (Figure 1E; Figure 2A).

### Analysis of phylogenetic and evolution of gene family

We identified 35,589 orthologous gene families between Sichuan taimen and nine other species, including 481 gene families specific to the Sichuan taimen and 342 single-copy orthologues. The results of the phylogenetic study showed that, as an ancient branch of Salmoninae, the divergence time between huchonids (*Hucho* spp. and *Brachymystax* spp.) and other Salmoninae fishes was estimated at approximately 33.29 Mya (Million years ago)(Figure 3; Supplementary Figure S4), which is consistent with previous studies (Horreo, 2017; Lecaudey et al., 2018). The divergence time estimated by PAML was 15.81 Mya between the taimen and its sister group, lenok (*Brachymystax* spp.)(Figure 3).

**Figure 3.**
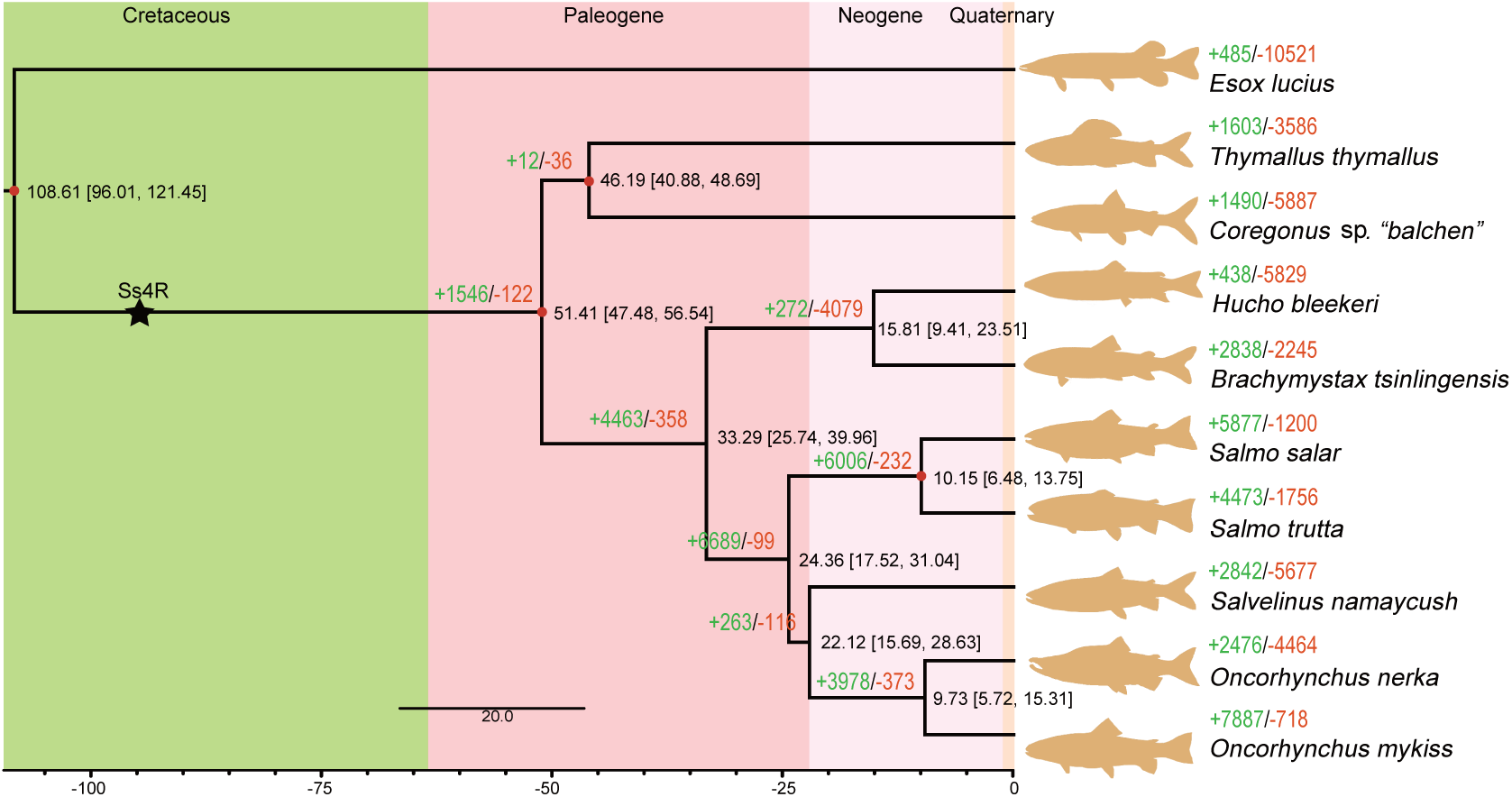
Maximum likelihood phylogenetic tree constructed based on single-copy orthologous genes. Fossil calibration points used by MCMC are marked in red. Black numbers next to each nodes are the divergence times (Mya) and the 95% confidential intervals estimated by PAML. The green positive and red negative integers indicate the number of genes at expanded or contracted gene families at that node, respectively (*P* < 0.05). Ss4R: salmonid-specific fourth vertebrate whole-genome duplication.

We identified 235 significant expanded genes in the Sichuan taimen genome (*P* < 0.01). Gene Ontology analysis showed the highest enrichment in nucleosome (GO:0000786) and chromosome (GO:0005694) categories (Supplementary Table S8). Similarly, a large number of histone genes were found in the expanded gene set, which we speculated might be influenced by the expansion of repetitive sequences. Besides, the GO term fin development (GO:0033333) was annotated, implying that the Sichuan taimen might have evolved enhanced swimming or predation abilities (Supplementary Table S8). KEGG pathway analysis showed enrichment in the Hedgehog signaling pathway (dre04340) and Insulin signaling pathway (dre04910), suggesting changes in embryonic development, energy metabolism and growth, which might be related to its special body size and habits (Supplementary Table S9).

### Ancestral chromosome status

To evaluate the chromosome evolution of Sichuan taimen, genomic synteny blocks were recognized against the Northern pike (*Esox lucius*), which serves as an appropriate outgroup preceding the Ss4R. A total of 12,077 orthologs (1:2 or 1:1 for *E. lucius* to *H. bleekeri*) were identified, resulting in the linkage of 80 homologous blocks covering 90.89% of the length of the 44 Sichuan taimen chromosomes (Supplementary Table S10). We recovered Sichuan taimen chromosome equivalents for all Northern pike chromosomes, with each Northern pike chromosome corresponding to two Sichuan taimen homologues (Figure 4A). The results showed that half of the Sichuan taimen chromosomes (26/44) mapped completely to individual Northern pike chromosomes (Figure 4A).

**Figure 4.**
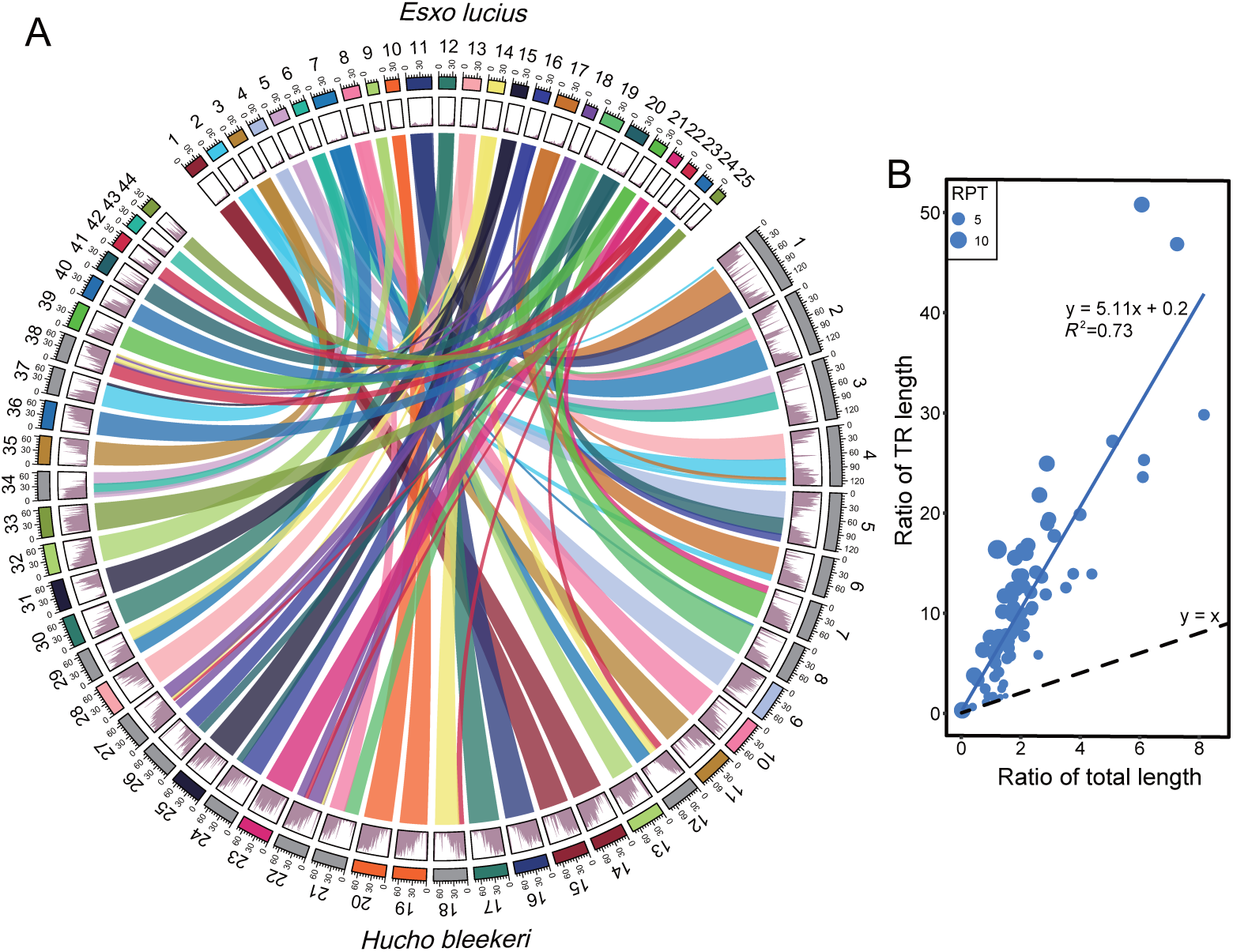
Chromosome evolution of *H. bleekeri*. A: Circos plot of homologous blocks and TR ratio in *E. lucius* and *H. bleekeri*. From the outside to the inside were chromosome length, TR ratio, and homologous block connections. B: TR expansion in taimen genome. The abscissa and ordinate were the ratio of the total length and TR length in homologous blocks between Sichuan taimen and Northern pike, respectively. The size of the circles was expressed as the ratio of the proportion of TR (RPT) in homologous blocks between them.

The total chromosome lengths were 3.32 Gb and 0.92 Gb for Sichuan taimen and Northern pike, respectively. However, the length of the Sichuan taimen genome far exceeded what would be expected from whole-genome duplication alone. The expansion of TR in the Sichuan taimen genome was also evident in the circos plot (Figure 4A). We further evaluated the TR expansion in the Sichuan taimen by comparing the ratio of total length to TR length in the 80 homologous blocks. The results revealed that all blocks showed TR expansion in the Sichuan taimen genome, with the TR length ratio increasing proportionally with total length growth (Figure 4B). Linear regression analysis indicated that the TR ratio in the Sichuan taimen genome was approximately fivefold higher than that of the Northern pike (Figure 4B).

### Demographic history and divergence for two taimen species

The geographical distribution pattern of the other landlocked huchonid fish, lenok, is similar to that of taimen. Both are found in the Qinling Mountain regions and the Heilongjiang River, and the competitive relationship is existed between them (Galland et al., 2021; Kaus et al., 2019). To reveal the divergence time between the two taimen species, one Siberia taimen (*H. taimen*) sample were collected from the Heilongjiang River (Figure 5A). A total of 44,120 genes were captured from the Siberia taimen genome, and we identified 4,860 1:1 orthologous genes among the four species. The result of BEAST showed that two taimen species separated at 2.29 Mya (Figure 5B), which was similar to the previous inference (Kucinski & Fopp-Bayat, 2022).

**Figure 5.**
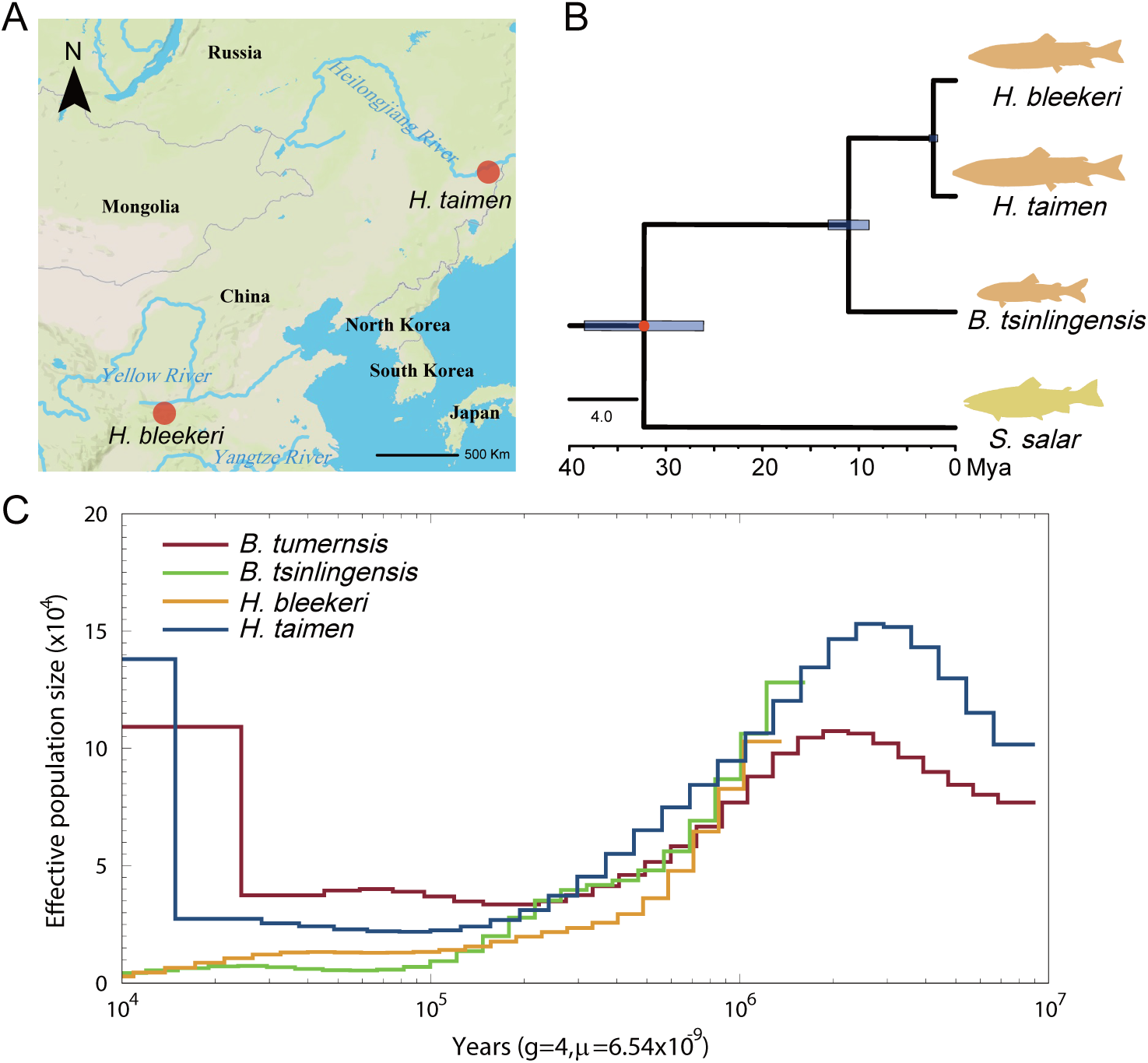
Species divergence history between *H. taimen* and *H. bleekeri*. A: Map of the two taimen species distribution. B: Phylogeny and divergence time of huchonid. 95% confidence interval was labeled as blue frame. Fossil calibration points were marked in red. C: Demographic history of two Heilongjiang sample (*H. taimen* and *B.tumernsis*) and two Qinling sample (*H. bleekeri* and *B.tsinlingensis*). The generation (g) time was set at 4 years and the neutral mutation rate per generation (μ) was set at 6.54 × 10-9.

To illustrate the trends in historical effective population sizes (*N*_e_) changes of taimen and lenok in Shaanxi and Heilongjiang, four resequencing samples from these regions were used for joint analysis. The results showed that the histories of the four species trace back to different times: ∼9 Mya for the two Heilongjiang samples (*H. taimen* and *B.tumernsis*), and ∼1 Mya for the two Shaanxi samples (*H. bleekeri* and *B. tsinlingensis*)(Figure 5C). *H. taimen* had a higher *N*_e_ level than *H. bleekeri* throughout their entire history (Figure 5C). The *N*_e_ curve suggested that a large-scale population expansion event occurred around 2 Mya in both *H. taimen* and *B.tumernsis*, followed by a continuous decline in population size over millions of years (Figure 5C). In summary, the Heilongjiang populations had higher *N*_e_ level than the Shaanxi populations since ∼200 Kya. Although the biomass of Sichuan taimen is currently smaller than that of Qinling lenok, there might have been times in history when the opposite was true (Figure 5C), which might reflect the competitive dynamics between them.

## DISCUSSION

### Complete genome assembly under high tandem repeat ratio

The combination of HiFi and Hi-C reads allowed us to achieve the chromosome-level assembly of the largest salmonid genome to date. The final size of the Sichuan taimen genome was approximately 3.45 Gb, which exceeds the size of the Atlantic salmon genome (∼2.97 Gb)(Lien et al., 2016). The chromosome count of the Sichuan taimen genome was 44, which approached to other salmonids such as *Salvelinus namaycush* (42), *B. tsinlingensis* (41), and *Salmo trutta* (40)(Supplementary Table S11). The assembled genome recovered 95.20% complete BUSCO genes, with a relatively high percentage of duplicated BUSCOs (46.20%), similar to its sister group *B. tsinlingensis* (Smith et al., 2022). Additionally, the chromosomal-level gene synteny implied the integrity of the genome assembly and annotation.

The genome assembly process and outcome were affected by the ratio of repetitive sequences (Tørresen et al., 2017; Treangen & Salzberg, 2011). The highly accurate reads produced by CCS technology improved both the process and quality of the assembly, especially inrepetitive sequences regions (Logsdon et al., 2020; Wenger et al., 2019). We found that the assembled Sichuan taimen genome contains a high proportion of repetitive sequences, with nearly half of the genome covered by tandem repeat sequences. This may contribute to the increased genome size and pose challenges for genome assembly. Fortunately, most of the TR regions were covered and assembled by the HiFi reads, demonstrating the advantage of long reads. However, some long TRs accounted for a relatively large proportion of the total genome length, causing gaps around them. Generally, the number of gaps in the assembled Sichuan taimen genome was similar to or even smaller than other salmonid genomes, except for GCF_013265735.2, indicating that our assembly was not severely disturbed by TRs (Supplementary Table S11).

Long repetitive regions can affect gene annotation, leading to the appearance of false and redundant genes (Lang et al., 2020). In our preliminary gene annotation, we found many tandem pseudogenes in long TR regions, which interfered with subsequent analysis. Despite the presence of TRs in specific protein domains (Simon & Hancock, 2009), each of these pseudogenes usually had only one exon according to our manual inspection. Therefore, we removed these single-exon genes located within repetitive sequence regions based on expression levels (TPM<1). In parallel, we blasted these pseudogenes to confirm their absence in other species.

### Phylogenetic relationships of Salmoninae based on single-copy orthologous genes

Salmonidae contains three subfamilies: Coregoninae, Thymallinae, and Salmoninae. However, their evolutionary relationships remains uncertain (Dysin et al., 2022). Lineage-specific rediploidization, incomplete lineage sorting, hybridization, and introgression have resulted in a complex and disordered phylogenetic relationship within Salmonidae (Campbell et al., 2020; Lecaudey et al., 2018; Robertson et al., 2017). Some studies suggest that Thymallinae is the sister group of Salmoninae (Crête-Lafrenière et al., 2012; Mckay et al., 2004; Wang et al., 2011), while other studies place Thymallinae at the root of the Salmonidae phylogenetic tree (Horreo, 2017; Wang et al., 2022). Based on the phylogenetic tree constructed using the orthologous genes, we suggest that Coregoninae and Thymallinae have a sister relationship, supported by 97% bootstraps (Supplementary Figure S4). This is consistent with other trees based on orthologues (Macqueen & Johnston, 2014; Robertson et al., 2017), mitochondrial genome (Campbell et al., 2013; Shedko et al., 2013), and ultra-conserved elements (Campbell et al., 2020). The inferred topology within Salmoninae aligns with other studies, with the genera *Hucho* and *Brachymystax* forming the ancient branch of the subfamily, constituting a sister group to the genera *Salmo, Salvelinus*, *and Oncorhynchus* (Crespi & Fulton, 2004; Kucinski & Fopp-Bayat, 2022). The divergence time between taimen and lenok was estimated at 9.41 to 23.51 Mya, which aligns with previous results based on mitochondria (Kucinski & Fopp-Bayat, 2022), and coincides with the Miocene Climate Optimum (MCO) at 15-18 Mya (Böhme, 2003; Sun & Zhang, 2008).

### Effective population size changes and the urgency of conservation

All species of the genera *Hucho* and *Brachymystax* are endangered and protected by Chinese government law (Sung et al., 1998). The results of PSMC revealed that the effective populations size of four *Hucho* and *Brachymystax* species have declined over the past million years, following the separation of *H. bleekeri* and *H. taimen*. Similar patterns of *N*_e_ curves were observed in species with the same geographical distribution. The curves of *H. bleekeri* and *B. tsinlingensis* in Shaanxi showed a close coalescent endpoint, we speculated that these two salmonids shared the same immigration route during the glacial period and were retained in low-latitude areas during the interglacial period. The Sichuan taimen had a lower *N*_e_ level than Siberia taimen in our calculation result, and current resource surveys indicate that Sichuan taimen are very rare in the Qinling Mountains, having disappeared from most of their original distribution areas (Shen et al., 2006; Wang et al., 2016). Consequently, more effective conservation measures should be implemented to protect the Sichuan taimen.

## CONCLUSIONS

This study provides a comprehensive genome assembly of the Sichuan taimen, uncovering a novel tandem repeat sequences expansion phenomenon and offering valuable insights into the evolutionary history of this species. The detailed analysis of phylogenetic relationships and effective population size changes highlights the historical population dynamics of the Sichuan taimen and its relatives. The assembled genome serves as an essential resource for future research on the evolutionary biology and conservation genetics of salmonids. As the Sichuan taimen remains critically endangered, this genomic resource provides a foundation for developing effective conservation measures to ensure the survival of this ancient and unique species.

## AUTHOR CONTRIBUTIONS

Xin-miao Zhang, Li-xin Wang and Yu Jiang conceived and supervised the project. Dong-mei Xiong, and Sheng-long Jian the samples. Xin-miao Zhang, and Dong-mei Xiong carried out the bioinformatic analyses. Xin-miao Zhang wrote the manuscript. Sheng-long Jian, Yu Jiang and Li-xin Wang revised the manuscript. All authors have read and approved the final manuscript.

## ACKNOWLEDGEMENTS

This study was financially supported by the National Natural Science Foundation of China (32273138) and the latitudinal project foundation of China Power Construction Group Beijing Survey, Design and Research Institute Co., Ltd. (K4050422184). We thank the High-Performance Computing Platform of Northwest A&F University and the National Supercomputing Center in Xi’an and Hefei for providing computing resources.

## SUPPLEMENTARY FIGURE AND TABLE LEGEND

Supplementary Figure S1. Frequency distribution map in 17-mer.

Supplementary Figure S2. Hi-C interaction heatmap of Hucho bleekeri genome between 44 chromosomes and unmapped scaffolds.

Supplementary Figure S3. Gene hotspots of the hucho genome.

Supplementary Figure S4. Phylogenetic trees under 1,000 bootstraps.

Supplementary Table S1. Genome size estimation table at k=17.

Supplementary Table S2. Summarized benchmarking in BUSCO notation.

Supplementary Table S3. Summary of TE in 10 salmonids (%).

Supplementary Table S4. Summary of TE in 10 salmonids (length).

Supplementary Table S5. Statistics of TR for 10 salmonids.

Supplementary Table S6. Summary of STRs’ unit and total length.

Supplementary Table S7. Genomic synteny of 44 homeologous blocks.

Supplementary Table S8. The Gene Ontology annotation of the 235 positive selected genes.

Supplementary Table S9. The KEGG pathway annotation of the 235 positive selected genes.

Supplementary Table S10. Genomic synteny of 80 homeologous blocks between Northern pike (luc) and Sichuan taimen.

Supplementary Table S11. Summary of genome assembly results of 10 salmonids.

